# Synaptic connectivity and electrophysiological properties of the nucleus of the lateral olfactory tract

**DOI:** 10.1101/2023.12.31.573522

**Authors:** Sapir Penker, Naheel Lawabny, Aya Dhamshy, Tamar Licht, Dan Rokni

**Author notes:** **Correspondence:** Tamar Licht –, Dan Rokni –.

## Abstract

The sense of smell is tightly linked to emotions, a link that is thought to rely on the direct synaptic connections between the olfactory bulb and nuclei of the amygdala. A small number of amygdaloid nuclei are the recipients of such direct input from the olfactory bulb and their unique functions are not known. Among them, the nucleus of the lateral olfactory tract (NLOT) is unique in its developmental history and gene expression. NLOT has been very little studied and consequentially its function is unknown. Furthermore, formulation of informed hypotheses about NLOT function is at this stage limited by the lack of knowledge about its connectivity and physiological properties. Here, we used pseudo-rabies tracing methods to systematically reveal monosynaptic inputs into NLOT, and adeno-associated viruses to reveal NLOT projection targets. We found that the NLOT is interconnected with several olfactory brain regions and with the basolateral amygdala. Some of these connections were reciprocal, and some showed unique interhemispheric patterns. We tested the excitable properties of NLOT neurons and the properties of each of the major synaptic inputs. We found that the NLOT receives powerful input from piriform cortex, tenia tecta, and the basolateral amygdala, but only very weak input from the olfactory bulb. When input crosses threshold, NLOT neurons respond with calcium-dependent bursts of action potentials. This integration of olfactory and amygdalar inputs suggests that NLOT plays a role in behaviors that combine smell and emotion, possibly assigning emotional value to odors.

**Significance statement:** Despite the well-known functional links between olfaction and emotions, the physiological properties of these links remain largely understudied. One major pathway by which olfactory and emotional signals interact, is via the nucleus of the lateral olfactory tract (NLOT). NLOT has been little studied and its function is yet unclear. The lack of physiological information hinders informed hypotheses. Here, we characterize the synaptic and intrinsic properties of NLOT neurons. We show that the NLOT receives converging olfactory and amygdalar inputs, and that NLOT neurons respond to input with high-rate bursts of action potentials. This suggests that the NLOT, that harbors ∼2500 cells, encodes a low-dimensional signal that is of high importance. We hypothesize that the NLOT assigns emotional value to odors.

## Introduction

The intimate links between olfaction and emotions are thought to rely on the direct input from early olfactory processing centers to the amygdala (Herz, 2005; Soudry et al., 2011; Kadohisa, 2013; Kontaris et al., 2020). However, there are several pathways projecting olfactory signals to the amygdala and many of them remain poorly analyzed. The nucleus of the lateral olfactory tract (NLOT) is a major hub that is positioned at the caudal end of the lateral olfactory tract (LOT) and is interconnected with several olfactory brain regions as well as the basolateral amygdala (BLA) (Price, 1973; Broadwell, 1975; Scalia and Winans, 1975; Krettek and Price, 1978; Luskin and Price, 1983; Krabbe et al., 2019). NLOT is an amygdaloid nucleus that is also considered part of olfactory cortex since it receives direct synaptic input from the olfactory bulb (OB) (Krettek and Price, 1978). It displays a unique gene expression pattern that distinguishes it from surrounding amygdalar nuclei (Hochgerner et al., 2023). It is perhaps the least studied olfactory cortical region. Its functional connectivity and physiological properties are unknown and its function is yet unclear.

NLOT is evolutionarily conserved and can be found in both mammalian and non-mammalian species (Turner et al., 1978; Deryckere et al., 2022; Metwalli et al., 2022). In mammalians, it is more prominent in macrosmatic species (Crosby and Schnitzlein, 1982), supporting the notion that it plays an important role in odor-guided behaviors. Several other findings suggest that NLOT plays an important role in odor-guided behaviors and particularly in linking olfactory and emotional functions. First, chemical ablation of NLOT was reported to cause severe deficits in odor-guided behaviors (Vaz et al., 2017). Second, odor responses in NLOT are strongly influenced by odor valence (Tanisumi et al., 2021), similarly to the olfactory tubercle (OT) (Gadziola et al., 2015; Millman and Murthy, 2020). Third, perturbation of NLOT activity affects social behavior in rats, that is known to rely on olfaction and be affected by emotional state (Hernández-Pérez et al., 2022). Lastly, NLOT neurons show marked transcriptomic changes following fear conditioning (Hochgerner et al., 2023). Given these findings, it stands to reason that NLOT plays an important role in linking odors to emotional experiences.

The input sources and output targets of NLOT have not been fully determined. Reported inputs include olfactory brain regions such as the OB and piriform cortex (Price, 1973; Scalia and Winans, 1975; Luskin and Price, 1983), and the basolateral amygdala (BLA, (Krettek and Price, 1978)). Reported outputs include projections to the OB, the anterior olfactory nucleus (AON), piriform cortex (PC), the OT, and the BLA (Santiago and Shammah-Lagnado, 2004; Krabbe et al., 2019). The relative synaptic efficacies of the various inputs into NLOT in driving its activity are unknown.

NLOT was estimated to contain about 20000 neurons in rats (Vaz et al., 2016). In mice, we estimate that it contains about 2500 neurons, less than one percent of the number of neurons in PC (Srinivasan and Stevens, 2017). This already suggests that NLOT may not be able to encode the full odor space, and probably operates on lower dimensional features. The cellular organization of NLOT is reminiscent of PC. It has three layers with layer 1 containing mostly neuropil. Its portion that is closest to the pial surface (layer 1a) contains the lateral olfactory tract (LOT) which consists of olfactory bulb mitral cell axons. Layer 2 contains most of NLOT neurons. However, unlike PC, layer 2 neurons are almost exclusively glutamatergic, pyramidal-like neurons (McDonald, 1983; Millhouse and Uemura-Sumi, 1985; Hernández-Pérez et al., 2022). These send their apical dendrites ventrally into layer 1 where they branch and form synaptic connections with olfactory bulb mitral cell axons. Layer 3 neurons are more diverse and less well characterized.

In this study, we developed a virus-based method to specifically infect NLOT neurons and used it to trace NLOT input sources and output targets. Additionally, we characterized the synaptic properties of the major inputs into the NLOT and the firing properties of NLOT neurons in acute slices.

## Methods

All procedures were performed using approved protocols in accordance with institutional (Hebrew University Institutional Animal Care and Use Committee) and national guidelines.

### Mice

For activation of olfactory bulb inputs, Tbet-Cre mice (Jackson Laboratory strain 024507 (Haddad et al., 2013)) were crossed with Ai32 mice (Jackson Laboratory strain 024109, (Madisen et al., 2012)). C57bl6 mice were used in all experiments in which ChR2 expression was driven virally. Rbp4-cre mice (MGI:4367067, (Gong et al., 2003)) were used to specifically express the rabies helper viruses in the NLOT. 2–6-month-old male and female mice were used for connectivity tracing experiments. For electrophysiology, AAVs carrying the channelrhodopsin (ChR2) gene were injected into 3–6-week-old males. Animals were kept in a regular specific pathogen–free housing conditions with a 12-hour light/ 12-hour dark cycle. Irradiated rodent food and water were given *ad libitum*.

### Virus injections

Mice were anesthetized with isoflurane, given analgesia (Meloxicam 5mg/kg) and were placed in a stereotactic device (Kopf). A small craniotomy was made over the desired injection site and viral solution was injected into the brain with a fine syringe (volume: 80-150 nl at a rate of 100 nL/min, MICRO-2T, WPI). The coordinates for the different brain regions are shown in Table 1.

**Table 1.** Coordinates used for virus injections.

| Brain region | Anterior-posterior | Medial-lateral | Depth (from pia) |
| --- | --- | --- | --- |
| Olfactory bulb | 4.7 | 1 | 1.5 |
| Dorsal tenia tecta / AON | 2.3 | 0.3 | 3.5 |
| aPC | 1.2 | 3 | 4 |
| OT / Islands of Calleja | 1.5 | 2 | 4 |
| NLOT | -0.5 | 2 | 5.5 |
| BLA | -1.5 | 3.1 | 4.5 |
| pPC | -2 | 3.6 | 4.5 |

The syringe was then slowly retracted, the craniotomy was sealed with bone wax, and the skin was sutured. A minimum of 3 weeks were allowed for all adeno-associated viruses (AAVs) to express. Pseudotyped-rabies were allowed 5-7 days to express. All adeno-associated viruses (AAVs) were made from AddGene plasmids by the ELSC vector core facility at the Hebrew University (EVCF). EnvA pseudotyped-G deleted rabies virus expressing GFP (EnvA-ΔG-RV-GFP) was purchased from the viral vector core of Kavli Institute for Systems Neuroscience, Norway. All viruses used in this study are listed in Table 2.

**Table 2.**
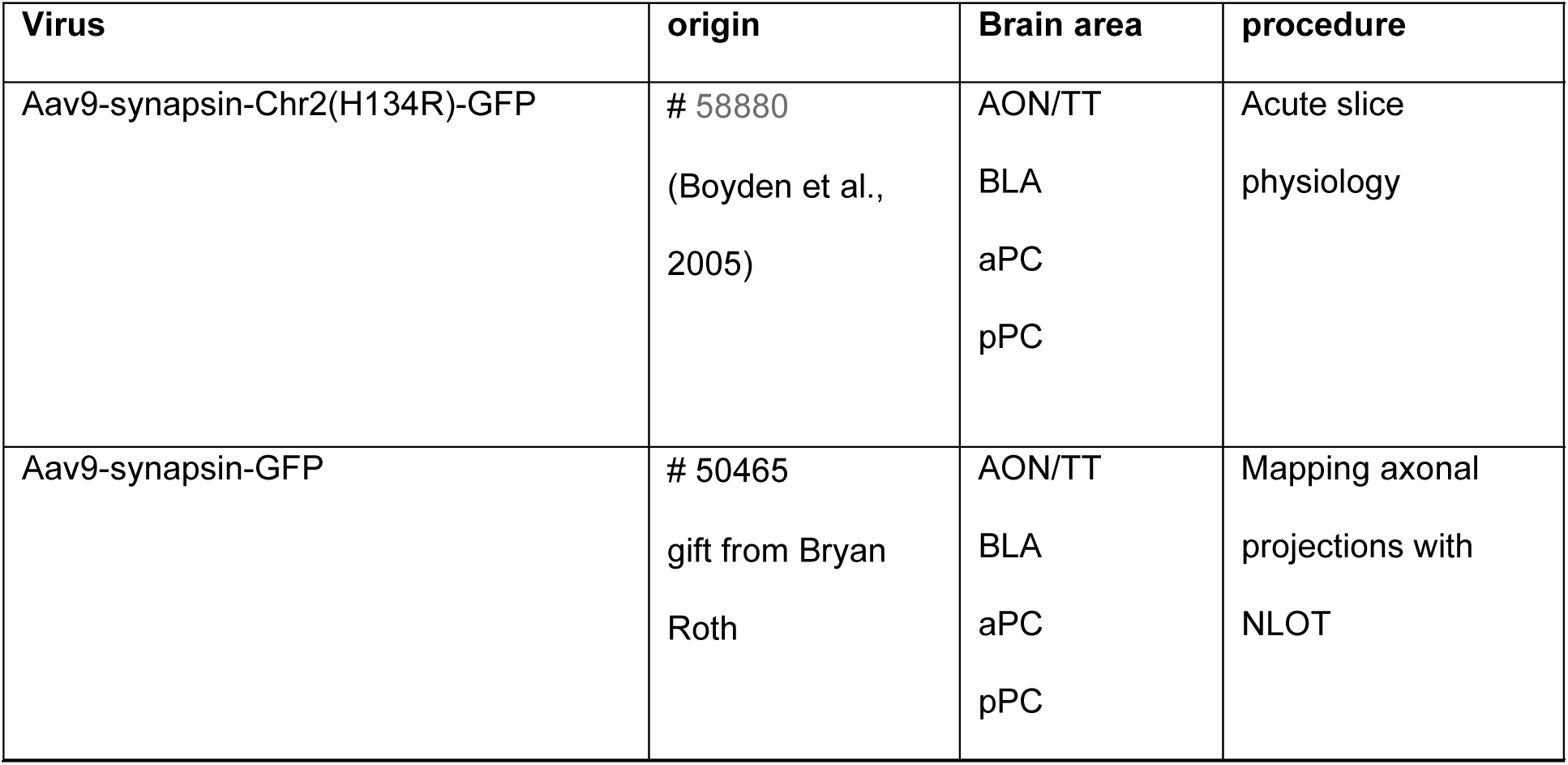
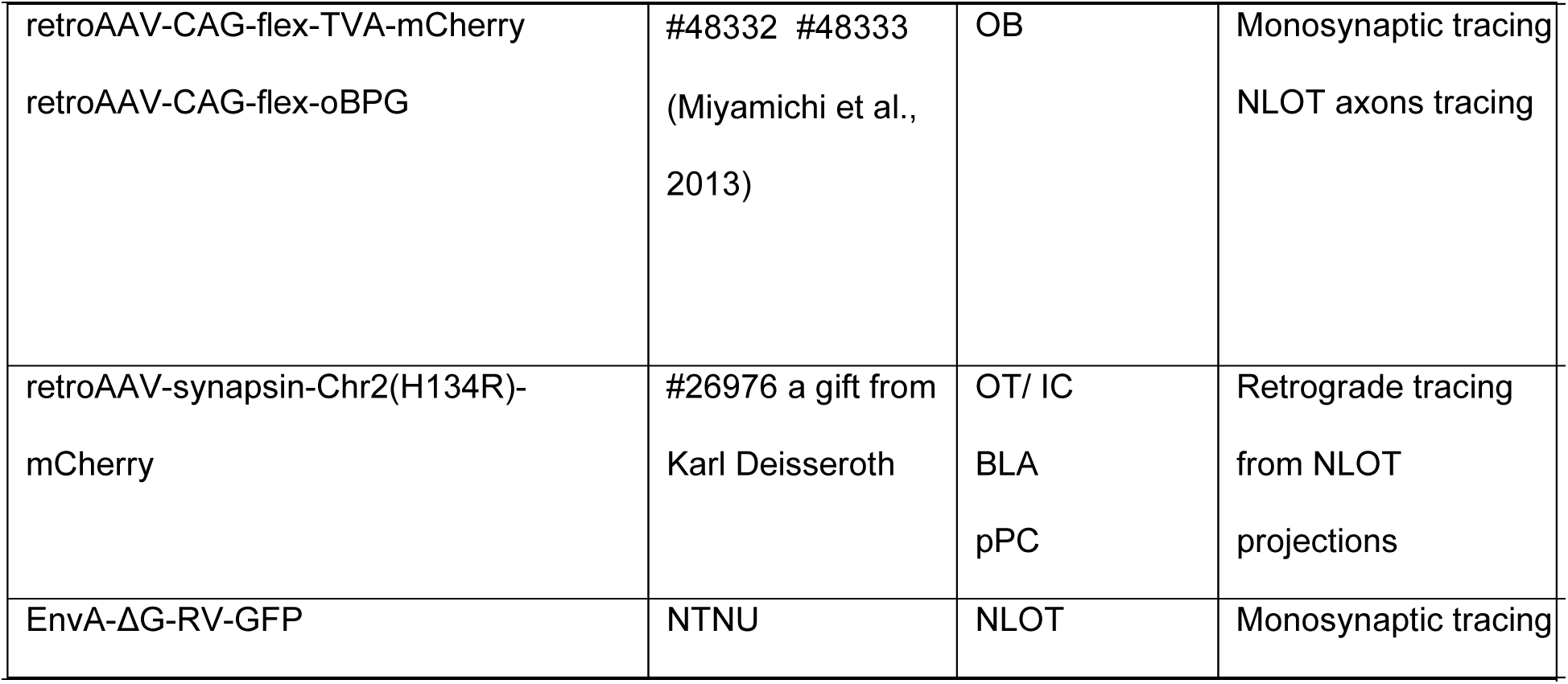
Viruses used in this study.

| Virus | origin | Brain area | procedure |
| --- | --- | --- | --- |
| Aav9-synapsin-Chr2(H134R)-GFP | # 58880<br>(Boyden et al., 2005) | AON/TT<br>BLA<br>aPC<br>pPC | Acute slice<br>physiology |
| Aav9-synapsin-GFP | # 50465<br>gift from Bryan Roth | AON/TT<br>BLA<br>aPC<br>pPC | Mapping axonal<br>projections with<br>NLOT |
| retroAAV-CAG-flex-TVA-mCherry<br>retroAAV-CAG-flex-oBPG | #48332 #48333<br>(Miyamichi et al.,<br>2013) | OB | Monosynaptic tracing<br>NLOT axons tracing |
| retroAAV-synapsin-Chr2(H134R)-<br>mCherry | #26976 a gift from<br>Karl Deisseroth | OT/ IC<br>BLA<br>pPC | Retrograde tracing<br>from NLOT<br>projections |
| EnvA-ΔG-RV-GFP | NTNU | NLOT | Monosynaptic tracing |
| **Table 2. Viruses used in this study** |  |  |  |

### Slice Electrophysiology

#### Slice preparation and solutions

Mice were anesthetized with 5% Isoflurane (RWD Vaporizer for isoflurane), decapitated, and 300 µm coronal slices were made in ice-cold solution containing (in mM: NaCl 126, KCl 2.5, MgCl_2_ 1.2, NaH_2_PO_4_ 1.4, CaCl_2_ 2.4, Glucose 11, NaHCO_3_ 25, Ascorbate 0.4, Sodium pyruvate 2, Kynurenic acid 2.5, NaOH 0.6) and equilibrated with 95% O2/5% CO2. Slices were transferred to a holding chamber containing (in mM: NaCl 126, KCl 2.5, MgCl_2_ 1.2, NaH_2_PO_4_ 1.4, CaCl_2_ 2.4, Glucose 11, NaHCO_3_ 25, Ascorbate 0.4, Sodium pyruvate 2) equilibrated with gases as above and let to rest for at least 30 min at room temperature. During recordings slices were continuously perfused with the same solution as in the holding chamber at 32° C. For voltage clamp experiments, patch pipettes (2 Mohm) were filled with (in mM: CsMS 128, HEPES-K 10, EGTA 1, MgCl2 1, NaCl 10, Mg-ATP 2, Na-GTP 0.3, at pH 7.2–7.4). For current clamp experiments the internal solution contained (in mM: D-Gluconic acid 130, EGTA 1, HEPES-K 10, KCL 20, MgCl_2_ 1, Mg-ATP 2, Na-GTP 0.3, at pH 7.2–7.4). In some of the experiments, the internal solution also contained biocytin (5.36 mM, B4261 Sigma-Aldrich) to allow post hoc visualization of cellular morphology. The following ionic channel and synaptic blockers were added to the ACSF and applied via the perfusion system: NiCl 200 µM (Sigma-Aldrich), TTX 1 µM (Alomone labs), APV 60 µM (Alomone labs), CNQX 10 µM (Alomone labs).

#### Data acquisition

Slices were visualized with an Olympus BX51 microscope mounted on a Luigs and Neumann shifting table. Current and voltage were recorded using a MultiClamp 700B amplifier (Molecular Devices), digitized with 16-bit resolution and sampled at 10KHz (Axon Digidata 1550B). PClamp (Molecular Devices) was used for data acquisition as well as controlling light stimulation and current and voltage-clamp protocols. An ultra-bright LED (470 nm, 1450mW, Prizmatix) that was mounted on the rear lamp housing of the microscope was for light stimulation. Light from the LED was passed through an Olympus U-M49002 filter set (ET470/40x, beam-splitter T495LP, emitter ET525/50m) and through the objective onto the focal plane. Light pulses ranged between 0.2 and 1 ms.

All analysis of electrophysiological data was performed using custom-written code in Matlab (Mathworks).

### Histology and microscopy

Brains or acute slices were fixed by immersion in 4% paraformaldehyde on ice for 12h. Brains were washed with PBS, and sectioned to 50-μm coronal sections by a vibratome (Leica). Staining was performed for rabies tracing experiments as described (Licht et al., 2023) with anti-GFP (1:400; Abcam RRID: AB_305643) and Alexa 488 anti-goat (RRID: AB_2336933) as secondary antibody. Acute slices with biocytin-filled cells were stained with Streptavidin-cy3 (Jackson Immunoresearch) diluted 1:400. Sections were mounted on glass slides, covered by mounting medium containing Dapi (SouthernBiotech), and coverslipped. Low-magnification images were acquired using Nikon SMZ-25 fluorescent stereoscope equipped with X1 and X2 objectives. Confocal microscopy was done using Olympus FV-1000 on 10X and 20X objectives and 1µm distance between confocal z-slices.

## Results

### Intrinsic properties of NLOT neurons

We first analyzed the excitability of layer-2 NLOT neurons in acute coronal slices from young adult mice (1-4 months old). NLOT is readily recognized under bright-field illumination allowing easy identification (Figure 1A). We performed whole cell recordings using the current clamp mode to study resting membrane parameters and firing properties (Figure 1B). NLOT neurons had a resting membrane potential of ∼-65 mV (−67 ± 5 mV, median ± MAD, n=22), and an input resistance of ∼ 180 MΩ (179 ± 88 MΩ, median ± MAD, n=22, Figure 1C). At rest, NLOT neurons were quiescent and typically required a depolarization of a little over 10 mV to reach action potential threshold (11.1 ± 3.9 mV, median ± MAD, n=22). When supra-threshold currents were injected, NLOT neurons fired bursts of action potentials (Figure 1B,D-F). These bursts had an average firing rate of over 30Hz and rode over a slower depolarization (Figure 1G). The slower depolarization was also evident (although less prominent) when action potentials were blocked with the Na^+^ channel blocker TTX (Figure 1D-E). Ni^2+^ blocked the slow depolarization, indicating that it is caused by a voltage-dependent Ca^2+^ current. Ni^2+^ also reduced the burstiness of NLOT neurons as indicated by its effect on action potential threshold, the number of spikes evoked by just suprathreshold currents, the firing rate within a burst, and the duration of the burst (Figure 1F-G). The prolonged firing when Ca^2+^ channels are blocked also suggests that bursts are terminated by Ca^2+^-activated K^+^ currents (Faber and Sah, 2002). These data suggest that NLOT neurons convey information by brief, high-rate, bursts of action potentials that are the result of an interplay between voltage-gated sodium and calcium channels.

**Figure 1:**
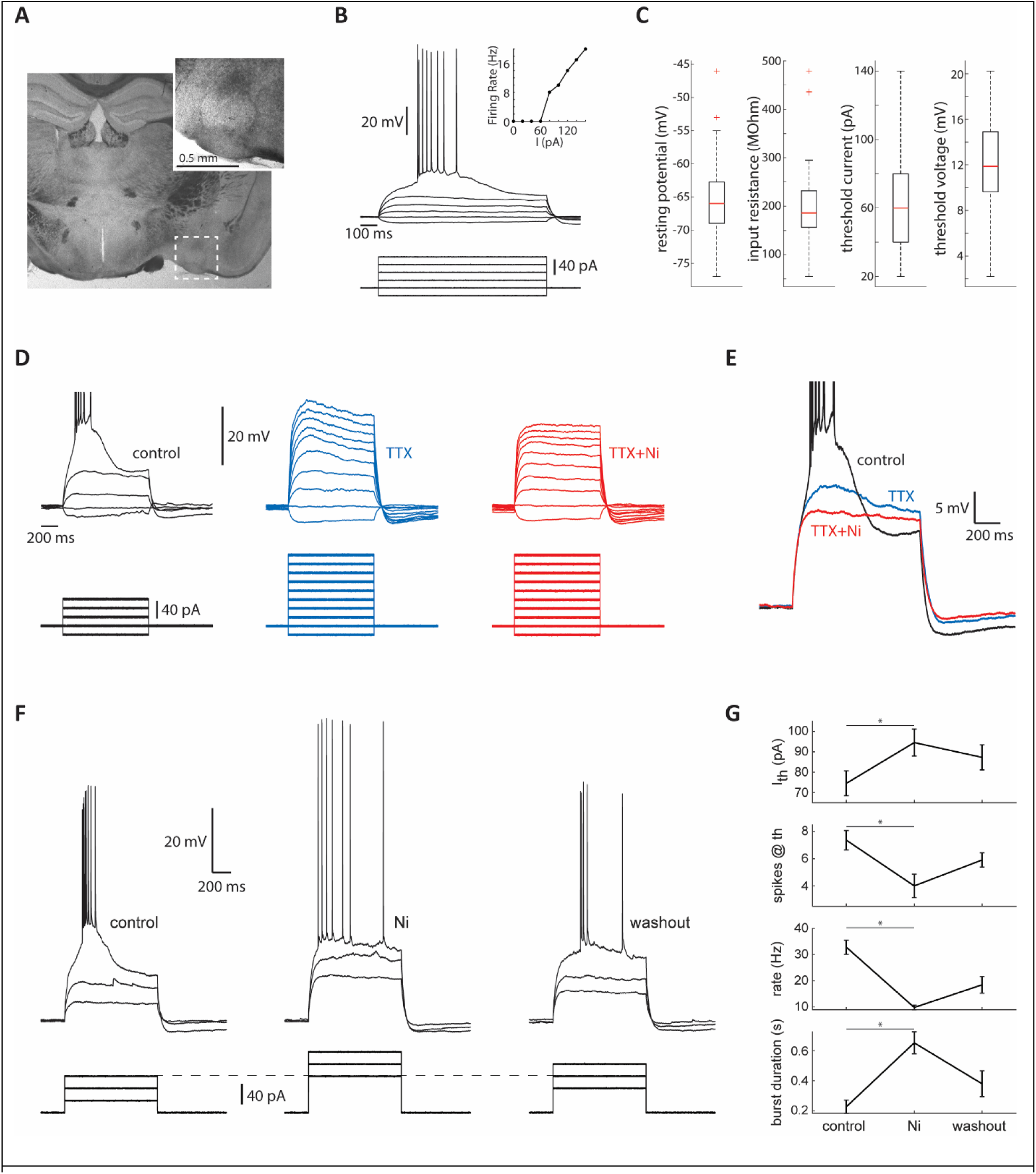
Calcium currents underly bursting in NLOT neurons. **A.** A coronal section showing the NLOT under bright-field illumination. **B.** The responses of a layer 2 neuron to current injections. Inset shows the F/I curve of this neuron. **C.** Box plots showing the distributions of resting membrane properties and spiking threshold of layer 2 neurons. **D.** The effect of the sodium channel blocker TTX (1 µM), and the calcium channel blocker Ni^2+^ (200 μM), on the responses to current injection in an example neuron. **E.** The superimposed responses of the neuron in D. Note that blocking action potentials with TTX reveals the contribution of a voltage gated calcium current. **F.** The effect of Ni2+ on bursting in an example neuron. **G.** quantification of the effect of Ni2+ on spiking threshold current, the number of spikes evoked by threshold current, the firing rate within the burst, and burst duration.

### NLOT connectivity

The connectivity of NLOT has not been systematically analyzed using modern tracing methods. To reveal its inputs and outputs we adopted virus-based tracing approaches. Because NLOT is small (∼0.5 mm) and ventral, direct virus injection almost always infects neighboring regions as well. Serendipitously, we found that NLOT is the only brain region that both projects to the main OB, and expresses Cre recombinase in the Rbp4-cre mouse line. Therefore, injections of Cre-dependent retrograde AAVs to the OB of Rbp4-Cre mice specifically infect NLOT neurons (Figure 2A,B). We used this approach to either label NLOT neurons and their projections and identify putative output targets, or to express helper viruses for monosynaptic rabies-based tracing of input sources.

**Figure 2:**
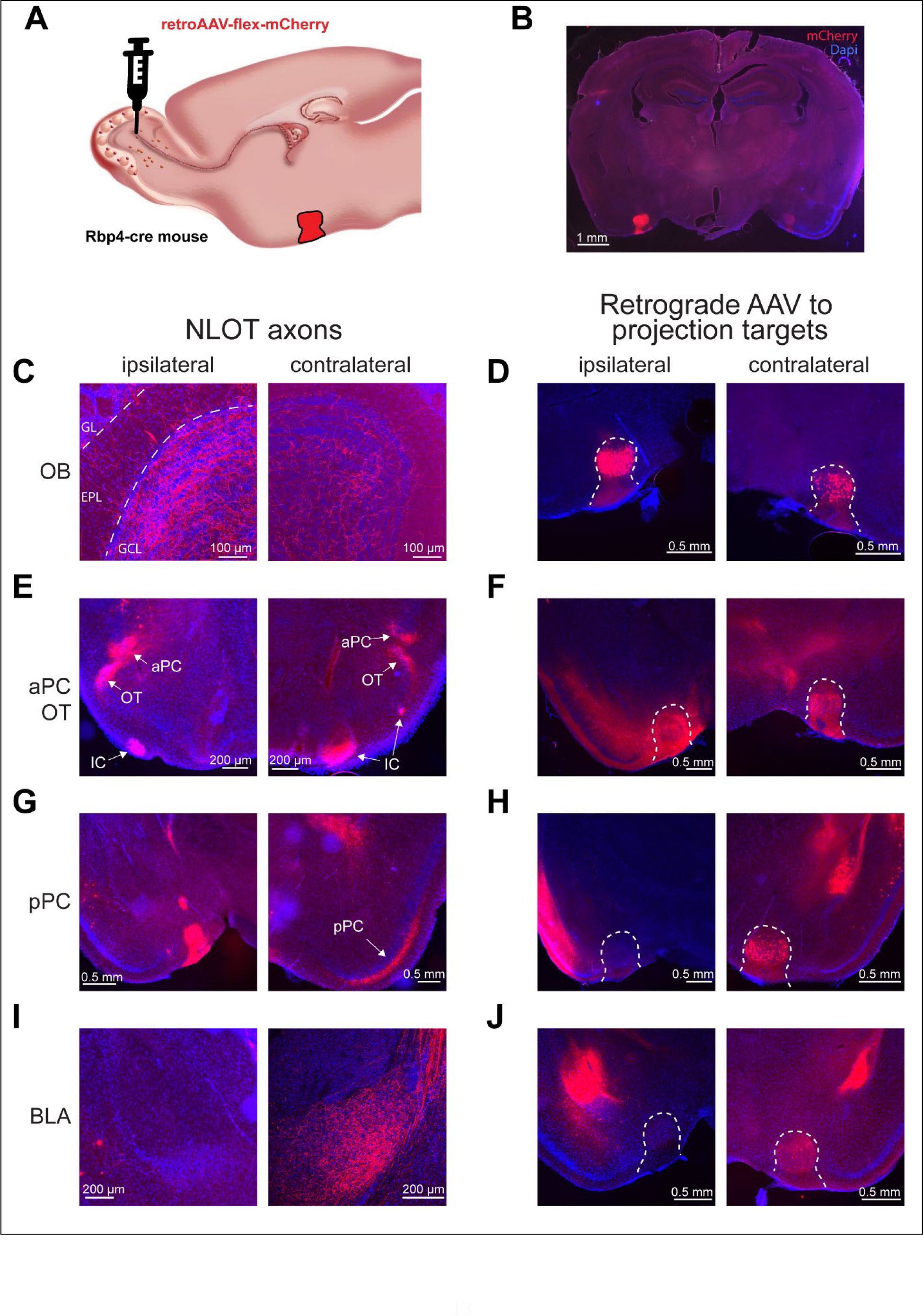
Axonal projections of NLOT neurons. **A.** Experimental schema. NLOT neurons were specifically labeled by a unilateral injection of retrograde AAV carrying the gene for mCherry into the OB. **B.** A coronal section showing the labeled NLOT. **C.** Sections of the ipsi- and contra-lateral OBs showing NLOT axons densely populating the granule cell layer (GCL) yet also reaching the external plexiform and glomerular layers (EPL and GL). **D.** The ipsi- and contra-lateral NLOTs labeled by unilateral retrograde AAV injection into the OB. **E.** NLOT axons within the aPC and the islands of Calleja (IC) in the OT. Note that NLOT axons specifically target the bordering regions of the aPC and OT. **F.** The ipsi- and contra-lateral NLOTs were labeled by retrograde AAV injection into the OT. **G.** NLOT projections in layer-1b of the contralateral, but not in the ipsilateral pPC. **H.** Retrograde AAV injection into pPC labeled the contralateral but not the ipsilateral NLOT. **I.** NLOT axonal projections within the contralateral BLA. No axons were evident in the ipsilateral BLA. **J.** Retrograde AAV injection into the BLA labeled the contralateral, but not the ipsilateral NLOT.

#### NLOT Output projections

We injected retrograde AAVs expressing Cre-dependent mCherry into one OB in each animal and examined the pattern of NLOT axonal projections across the brain’s two hemispheres. We found axonal projections in the OB, PC, the OT, and the BLA. We then complemented these experiments by Injecting retrograde AAVs expressing mCherry into these output targets and verifying that NLOT cells are labeled.

In the OB, labeling was densest within the granule cell layer but some axons crossed all the way to the glomerular layer (Figure 2C). This was true in both the ipsilateral and the contralateral OBs, but stronger labeling was evident in the ipsilateral side. Retrograde labeling from a single OB labeled neurons in the NLOT of both hemispheres but many more cells were labeled in the ipsilateral side (Figure 2D). The anterior piriform cortex (aPC) was mostly devoid of NLOT axons other than the ventromedial-most region bordering the OT. Within the OT axons were evident in the area bordering PC (presumably the anterolateral isolation of the CAP compartments (aiCAP) region that is the target of projections from a specific subpopulation of mitral cells (Hirata et al., 2019)), and within two lateral islands of Calleja (Figure 2E). This pattern was bilateral and was verified by unilateral retrograde AAV injection into the OT that labeled the NLOT in both hemispheres (Figure 2F). Projections to the posterior piriform cortex (pPC) were contralateral exclusively. These axons were evident in layer 1b suggesting that they may form synaptic connections close to the soma of pyramidal neurons and could potentially efficiently activate them (Figure 2G). Unilateral Injection of retrograde AAV into the pPC confirmed this cross-hemispheric connection by labeling NLOT cells only in the contralateral side (Figure 2H). NLOT projections to the BLA were also exclusively contralateral (Figure 2I), as was also confirmed by unilateral injection of retrograde AAV into the BLA (Figure 2J). We did not find any spatial organization of NLOT axons within the BLA.

To conclude, we show that the NLOT innervates the OB in both hemispheres (with more innervation in the ipsilateral OB), the aiCAP region and lateral Islands of Calleja in both hemispheres equally, and the pPC and BLA exclusively in the contralateral hemisphere. This is a rare example of exclusive cross-hemispheric connectivity involving two different brain areas.

#### Inputs into NLOT

To identify input sources into NLOT, we used the monosynaptic glycoprotein-deleted rabies virus tracing method. In order to selectively express the helper viruses in the NLOT, we injected Cre-dependent retrograde AAVs carrying the genes for TVA-mCherry and oPBG into the olfactory bulb of Rbp4-cre mice and allowed 4 weeks for the virus to be expressed in NLOT neurons. We then injected the modified rabies expressing GFP (rRb-EnvA-GFP) into NLOT (Figure 3A). 5-7 days later, we fixed the brains and made histological sections. As expected, we found starter cells (expressing both GFP and mCherry) only in the NLOT (Figure 3B).

**Figure 3.**
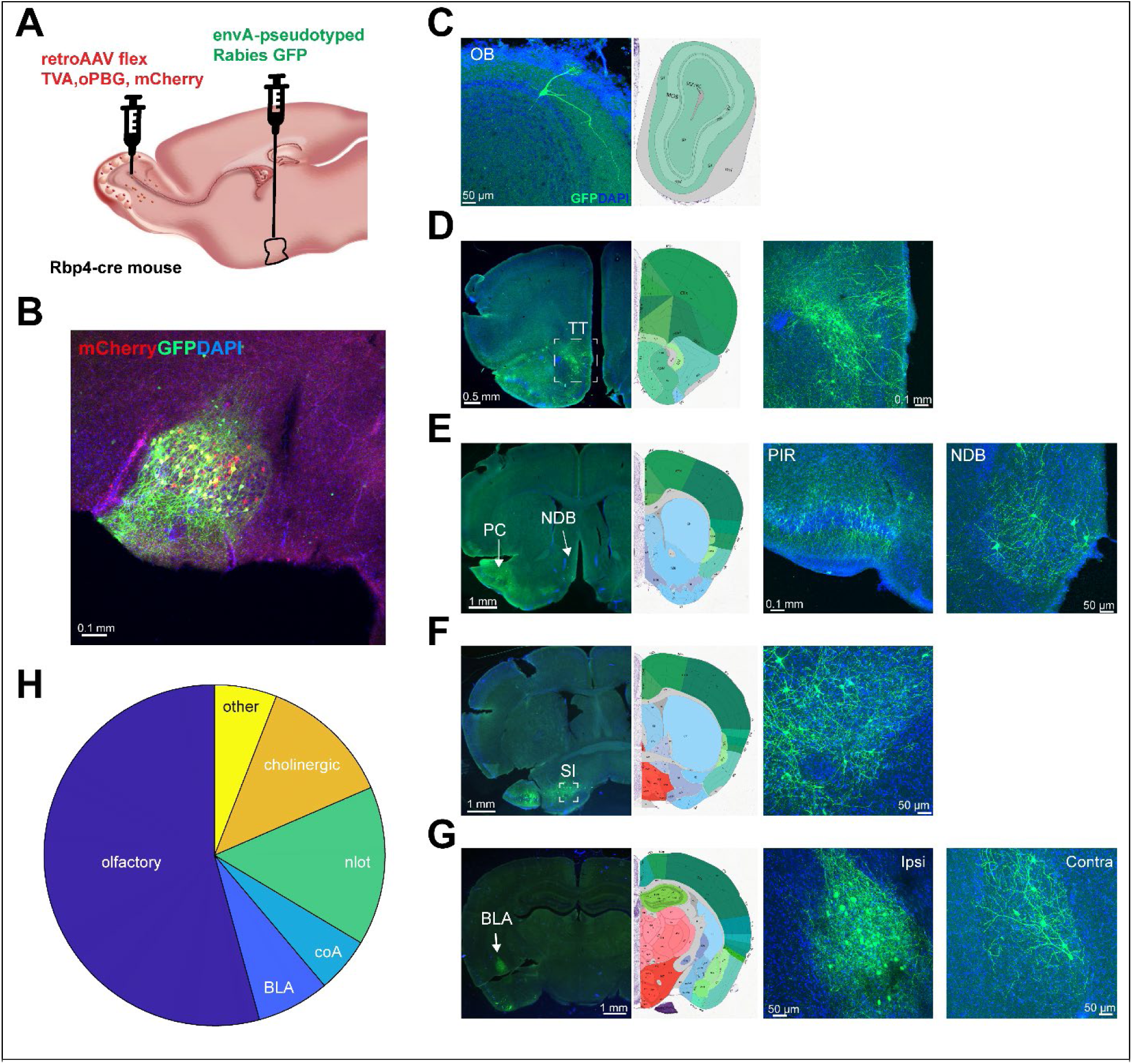
Inputs into the NLOT as revealed with g-deleted rabies monosynaptic tracing. **A.** Experimental scheme. Two retrograde AAVs, one carrying the genes for TVA and mCherry, and one carrying the gene for oPBG were injected unilaterally into the OB of Rbp4-Cre mice and allowed 4 weeks for expression. Then a pseudotyped rabies virus carrying the gene for GFP was injected into the ipsilateral NLOT. **B.** Fluorescently labeled cells in the ipsilateral NLOT. Yellow cells express both mCherry and GFP and are presumed to be starter cells. The occurrence of cells expressing only GFP indicates synaptic connectivity within the NLOT. **C-G.** Rabies-labeled neurons in the ipsilateral OB (C), ipsilateral tenia tecta (D), ipsilateral PC and NDB (E), ipsilateral SI (F), and ipsi- and contra-lateral BLA (G). **H.** The fraction of rabies-labeled cells in different regions of the brain. The olfactory category includes the OB, PC, AON, TT, and the OT. The NLOT and BLA categories include both hemispheres. The cholinergic category includes the SI and the NDB.

Afferent inputs, labeled with GFP were found in 7 main brain regions: The main OB, tenia tecta (TT), PC, BLA, several nuclei of the cortical amygdala (CoA), the substantia innominate (SI), the nucleus of the diagonal band (NDB). Notably, we also found GFP-expressing cells in the NLOT indicating local synaptic connectivity. Some GFP-expressing cells were also found in the contralateral NLOT. For complete details of all GFP-expressing cells that were mapped in these experiments, see supplementary table 1.

In the OB, all labelled cells were mitral cells (Figure 3C). Their soma was in the mitral cell layer, a single apical dendrite projected to the glomerular layer where it formed a tuft, and lateral dendrites transverse the external plexiform layer. All labeled cells were in the ipsilateral OB. TT is a brain region that has been little studied. It is divided into dorsal and ventral parts that differ in their connectivity and presumably also in their function. Most cells that were labeled by the rabies virus were in the dorsal TT (Figure 3D). Layer specificity of the labeled cells within TT suggests that specific subgroups of TT neurons, project to the NLOT. Here as well all labeled cells were in the ipsilateral hemisphere. Rabies-labeled neurons were found in both the aPC and pPC (Figure 3E). Mostly layer-2 but also some layer-3 neurons were labeled. Labeled cells were only found in the ipsilateral PC. Additionally, we found labeled cells in two cholinergic centers – the nucleus of the diagonal band (NDB, Figure 3E), and the Substantia Innominata (SI, Figure 3F). In both of these regions labeled cells were only found in the ipsilateral hemisphere. The BLA was the main brain region that provides bilateral input into the NLOT. We found labeled cells in both the ipsilateral and the contralateral BLA without any apparent spatial organization (Figure 3G). Functionally, most of the labeled cells were within brain regions that are part of the olfactory system with the other two major sources being amygdalar nuclei (including the NLOT itself) and cholinergic centers (Figure 3H). These experiments indicate that the NLOT integrates unilateral olfactory information and bilateral BLA information, and may be modulated by cholinergic centers.

#### Synaptic physiology of the NLOT

Trans-synaptic tracing does not inform us of the strength and functional significance of synaptic connections (Rogers and Beier, 2021). To analyze synaptic efficacy, we turned to slice physiology experiments. We specifically analyzed the synaptic properties of inputs from the OB, PC, AON/TT, and the BLA. These inputs represent the majority of olfactory and amygdalar inputs and their integration by the NLOT may underlie olfactory-emotional linking. Each input was analyzed separately by selective activation with channelrhodopsin2 (ChR2) while recording synaptic currents and potentials in NLOT layer 2 neurons. Synaptic currents were studied using a Cs-based internal solution and synaptic potentials using a K-based internal solution. Biocytin was added to the internal solution for recovering the cellular morphology of the recorded neurons. We found that the NLOT receives strong input from the PC, the BLA, and TT, but only very weak input from the OB. Below we describe the properties of each of these inputs in detail.

We first analyzed OB inputs using transgenic mice that express ChR2 in all OB output neurons (Figure 4A, Tbet-ChR2 mice were obtained by crossing Tbet-Cre mice with ai32 mice). The lateral olfactory tract (LOT), containing the axons of OB mitral cells, was evident in these mice in the ventral most part of layer 1 (Figure 4B-C). Layer 2 neuron dendrites projected ventrally and often branched and sent several distal branches into this ventral portion of layer 1 where they could form synaptic connections with mitral cell axons (Figure 4C). Brief light pulses (0.1-1 ms) evoked synaptic currents that were blocked by CNQX and APV, indicating that these are glutamatergic synapses (Figure 4D). OB-evoked EPSC amplitudes were extremely small (38 ± 8 pA, mean ± AEM, n=25 cells), and reversal potential was extremely depolarized (56 ± 12 mV, mean ± SEM, n=8 cells), probably reflecting, at least in part, the electrotonic distance of the synaptic connections from the soma (Figure 4D-F). When evoked in the current clamp mode, OB inputs were never sufficient to evoke action potentials on their own. Only when combined with current injection that brought cells close to their firing threshold were OB inputs able to elicit spikes (Figure 4G). However, two features may boost the efficacy of OB inputs. First, OB inputs could in some cases evoke regenerative responses that are below the threshold for somatic action potentials (Figure 4H). These were all-or-none in nature, as evident by the bimodal responses to repeated presentations of the same stimulus. Furthermore, these regenerative responses were more often elicited when the cell was depolarized. These features are suggestive of local dendritic spikes. Second, the OB input is facilitating as indicated by pairs of stimuli (Figure 4D, I & J). These data indicate that the OB is incapable of driving NLOT activity in itself, but may have an effect in combination with other inputs.

**Figure 4:**
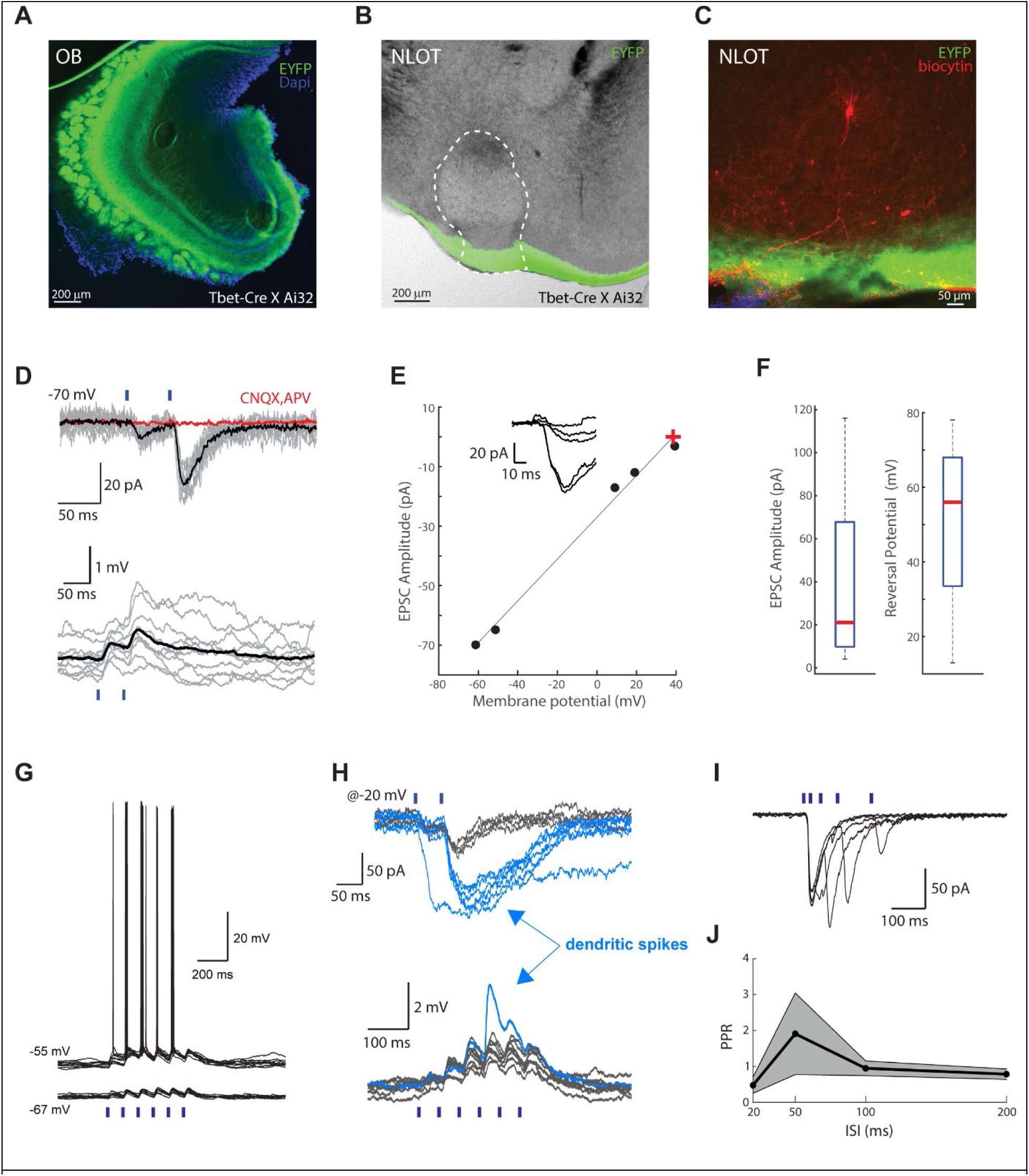
OB inputs are weak and insufficient to elicit NLOT firing. **A.** The expression of ChR2-EYFP in OB mitral and tufted cells. **B.** Fluorescent OB output axons (LOT) within layer 1 of the NLOT. **C.** A biocytin-labeled cell (red), and the LOT within layer 1. Note that the LOT contacts the dendrite at its distal end. **D.** Example LOT-evoked EPSCs (top) and EPSPs (bottom). Gray traces show individual trials and black lines show the average response across trials. Red line shows EPSC obtained in the presence of CNQX and APV. Blue ticks indicate the timing of light pulses. **E.** Voltage dependence of an example LOT-evoked EPSC. Red cross indicates reversal potential. **F.** Box plots showing the distribution of EPSC amplitudes (at −70 mV holding potential) and reversal potentials of LOT inputs. n=25 cells and 8 cells for the amplitude and reversal, respectively. **G.** LOT-evoked action potentials are only triggered when the cell is depolarized with current injection. **H.** LOT-evoked dendritic spikes recorded under voltage clamp (top) and current clamp (bottom) conditions (from two different cells). 10 repetitions of the same stimulation are shown for each. In the voltage clamp mode, the cell was held at −20 mV. Note the all-or-none nature of the responses that is indicative of a regenerative process (red). **I-J.** Paired-pulse dynamics of LOT inputs into NLOT neurons. **I.** Example data from one neuron. **J.** Paired-pulse ratio as a function of the inter-pulse interval averaged across cells (n=4). Note that LOT inputs are facilitating at intervals that are less than 100 ms.

Next, we describe the properties of inputs into NLOT from higher olfactory centers. We start with the PC. AAVs carrying the gene for ChR2 were stereotactically injected into the PC of young adult mice and were allowed 3-5 weeks’ time for expression (Figure 5A). We found no differences between the inputs from the anterior and posterior PC and combined them for the following description. PC axons were visible in the dorsal part of layer 1 and surrounding layer 3 where they could make contact with proximal dendrites of NLOT axons (Figure 5B). In line with that, brief light pulses evoked large responses in NLOT neurons (Figure 5C). EPSC amplitudes ranged between a few hundred pA and 2 nA (713 ± 106 pA, mean ± SEM, n=24) and the reversal potential of PC inputs was typical for glutamatergic synapses (9 ± 7 mV, mean ± SEM, n=11, Figure 5D-E). PC inputs showed strong paired pulse depression when the inter-stimulus interval was below 100 ms (Figure 5F-G). Importantly, PC inputs were sufficient to evoke action potential bursts similar to those seen with current injection (Figure 5H). Like with current injections, PC inputs typically evoked bursts of action potentials that rode over a slow depolarization lasting around 200 ms. Spike bursts tended to only occur in response to the earlier synaptic activations due to depression.

**Figure 5:**
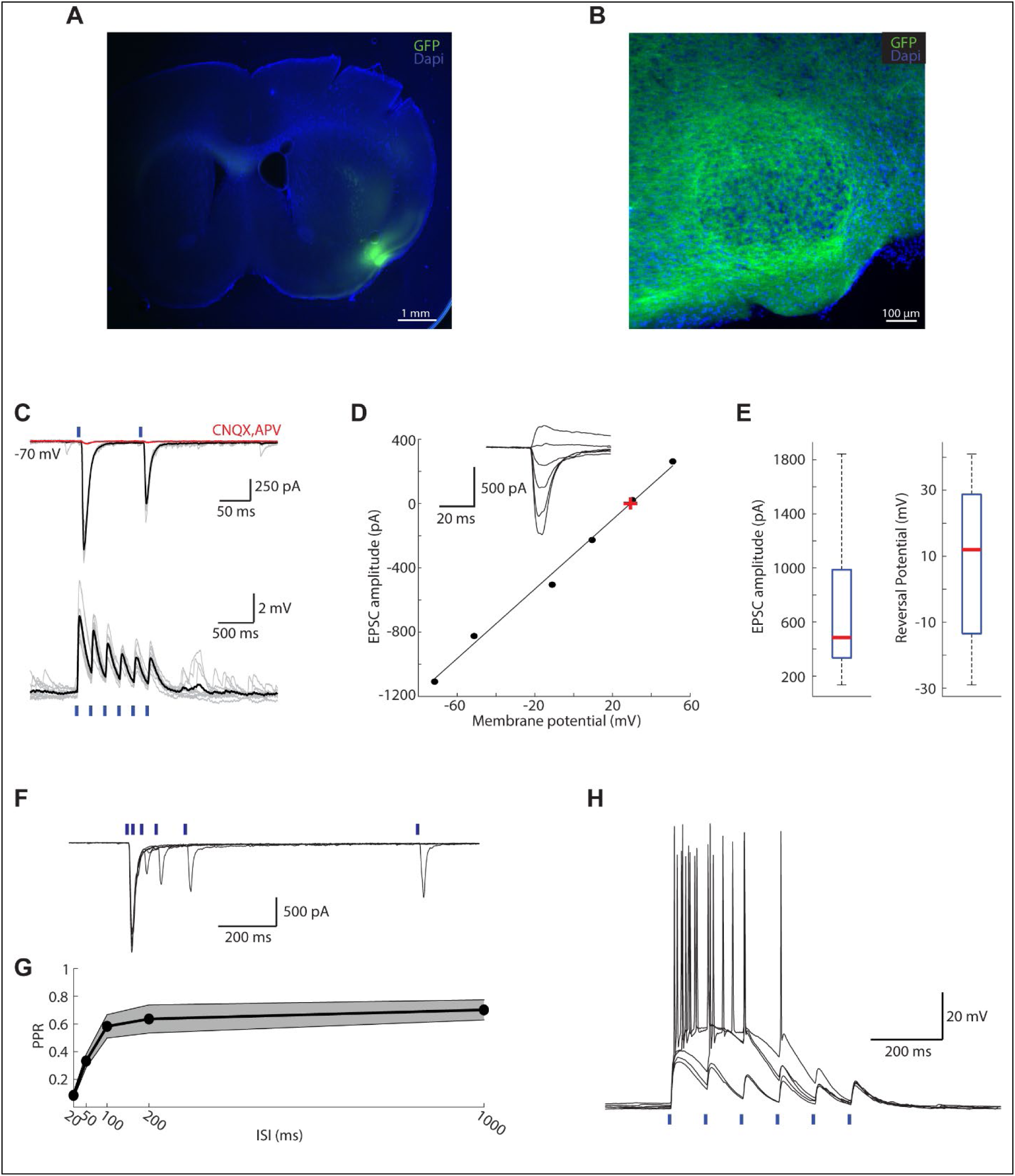
Piriform inputs into NLOT. **A.** A coronal section showing expression of ChR2-GFP in PC. **B.** A confocal image showing the spread of PC axons within the NLOT. **C.** Example PC-evoked EPSCs (top) and EPSPs (bottom). Gray traces show individual trials and black traces show their averages. Red trace shows the EPSC obtained in the presence of glutamatergic blockers. Blue ticks indicate the timing of light stimulation. **D.** Voltage dependence of an example PC-evoked EPSC. Red cross indicates reversal potential. **E.** Box plots showing the distributions of amplitudes and reversal potentials of PC-evoked EPSCs. n=24 and 11 cells for amplitude and reversal, respectively. **F-G.** Paired pulse dynamics of PC-evoked EPSCs. **F.** Example traces showing the EPSCs obtained with pairs of light pulses at variable intervals. **G.** The average paired pulse ratio as a function of the inter pulse interval. Shaded area shows the SEM (n=6 cells). **H.** PC-evoked action potential bursts. 5 trials are shown.

Next, we describe the properties of synaptic input from the TT/AON to the NLOT. TT/AON neurons were virally infected with ChR2 (Figure 6A). The TT is a narrow brain region and our injections often also infected the medial part of the anterior olfactory nucleus (AON). Injections that completely missed the TT and infected lateral parts of the AON failed to evoke synaptic input in NLOT neurons. Additionally, the rabies experiment showed many more labeled cells in the TT compared to the AON. However, as we could not isolate expression in either the medial AON or the TT, we treat these two possible inputs combined. TT/AON axons were visible within layer 2, suggesting that they contact NLOT neurons proximal to the soma (Figure 6B). Light pulses evoked EPSCs that were similar in their characteristics to the PC inputs but smaller in amplitude (403 ± 136 pA, mean ± SEM, n=13, Figure 6C). These inputs were reversibly blocked by glutamatergic blockers, reversed at normal glutamatergic reversal potential (11 ± 4 mV, mean ± SEM, n=13), and showed paired-pulse depression (Figure 6D-G). In the current clamp mode, light pulses readily evoked action potential bursts (Figure 6H).

**Figure 6:**
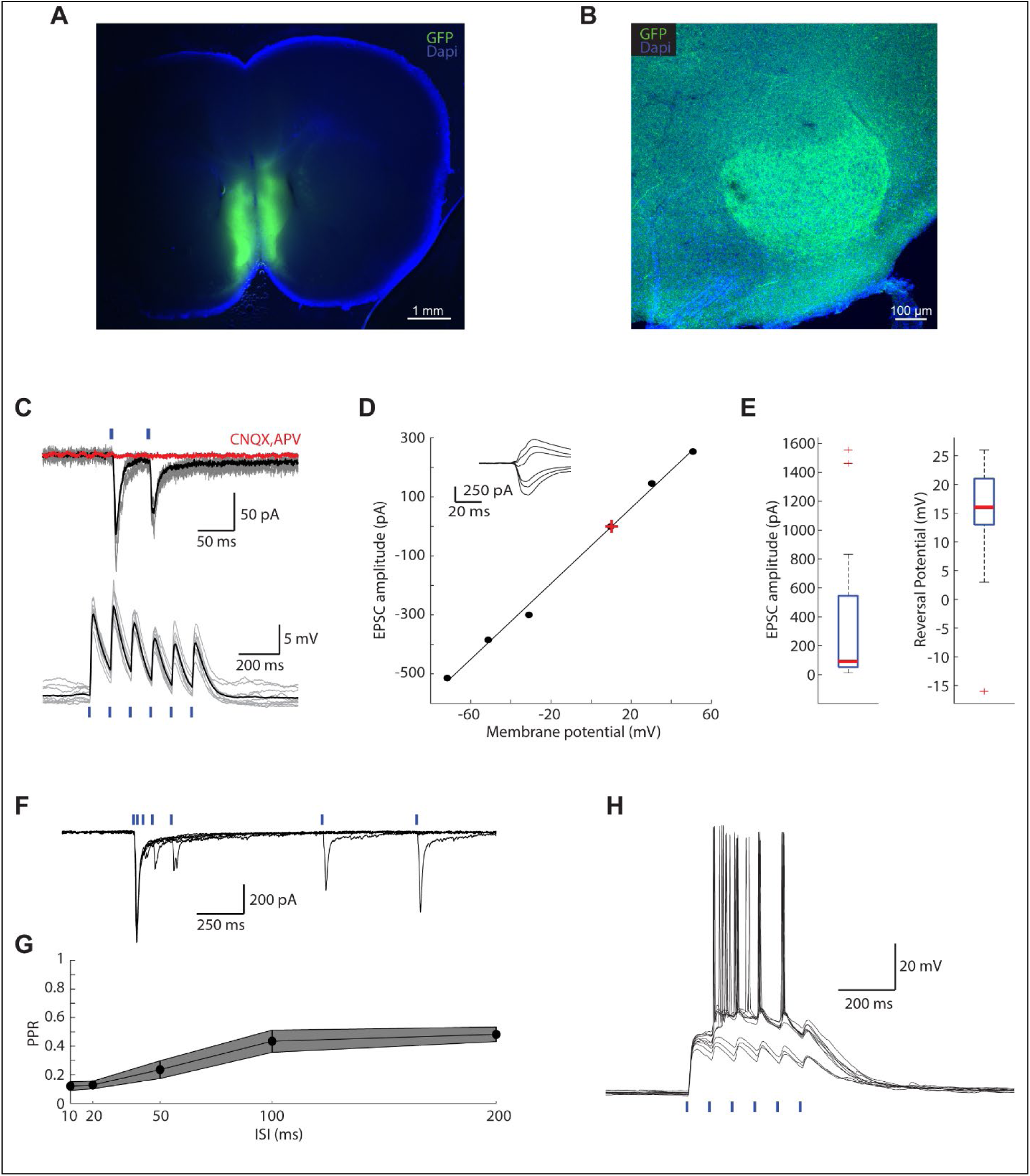
AON/TT inputs into NLOT. **A.** A coronal section showing the expression of ChR2-GFP in the medial-AON/TT. **B.** A confocal image showing the spread of AON/TT axons within NLOT. **C.** Example AON/TT-evoked EPSCs (top) and EPSPs (bottom). Gray traces show individual trials and black traces show their averages. Blue ticks mark the timing of light stimulation. **D.** Voltage dependence of EPSCs from one example cell. Black line is a linear fit. Red cross denotes reversal potential. **E.** Box plots showing the distributions of amplitudes and reversal potentials of AON/TT-evoked EPSCs (n=13 for both amplitude and reversal). **F-G.** Short-term synaptic plasticity of AON/TT inputs into NLOT. **F.** Responses from one example cell to paired stimulation with variable intervals. **G.** Average paired-pulse ratio as a function of inter-pulse interval. Shaded area shows the mean±SEM (n=9 cells). **H.** Example AON/TT-evoked action potential bursts. 10 repetitions of the same stimulation are shown.

These data indicate that of the olfactory inputs into the NLOT, third order regions (PC, AON/TT) that project more processed olfactory information are much more efficacious in activating NLOT neurons than the more peripheral OB.

Lastly, we analyze the input from the BLA to the NLOT. We expressed ChR2 in BLA neurons using viral injections (Figure 7A). Similar to the AON/TT axons, BLA axons were visible within layer 2 of the NLOT, suggesting that they innervate NLOT neurons at their proximal dendrites (Figure 7B). In line with that, light pulses evoked large and fast EPSCs (663 ± 100 pA, mean ± SEM, n=25 cells, Figure 7C). These were reversibly blocked by glutamatergic blockers (APV, CNQX), and reversed at −9 ± 12 mV (mean ± SEM, n=7, Figure 7D-E). Similar to the PC input, BLA synaptic input showed a marked pair-pulse depression at below 100 ms intervals (Figure 7F-G). BLA input was sufficient to evoke action potential bursts in NLOT neurons, but these tended to be limited to early stimuli within stimulation trains (Figure 7H).

**Figure 7:**
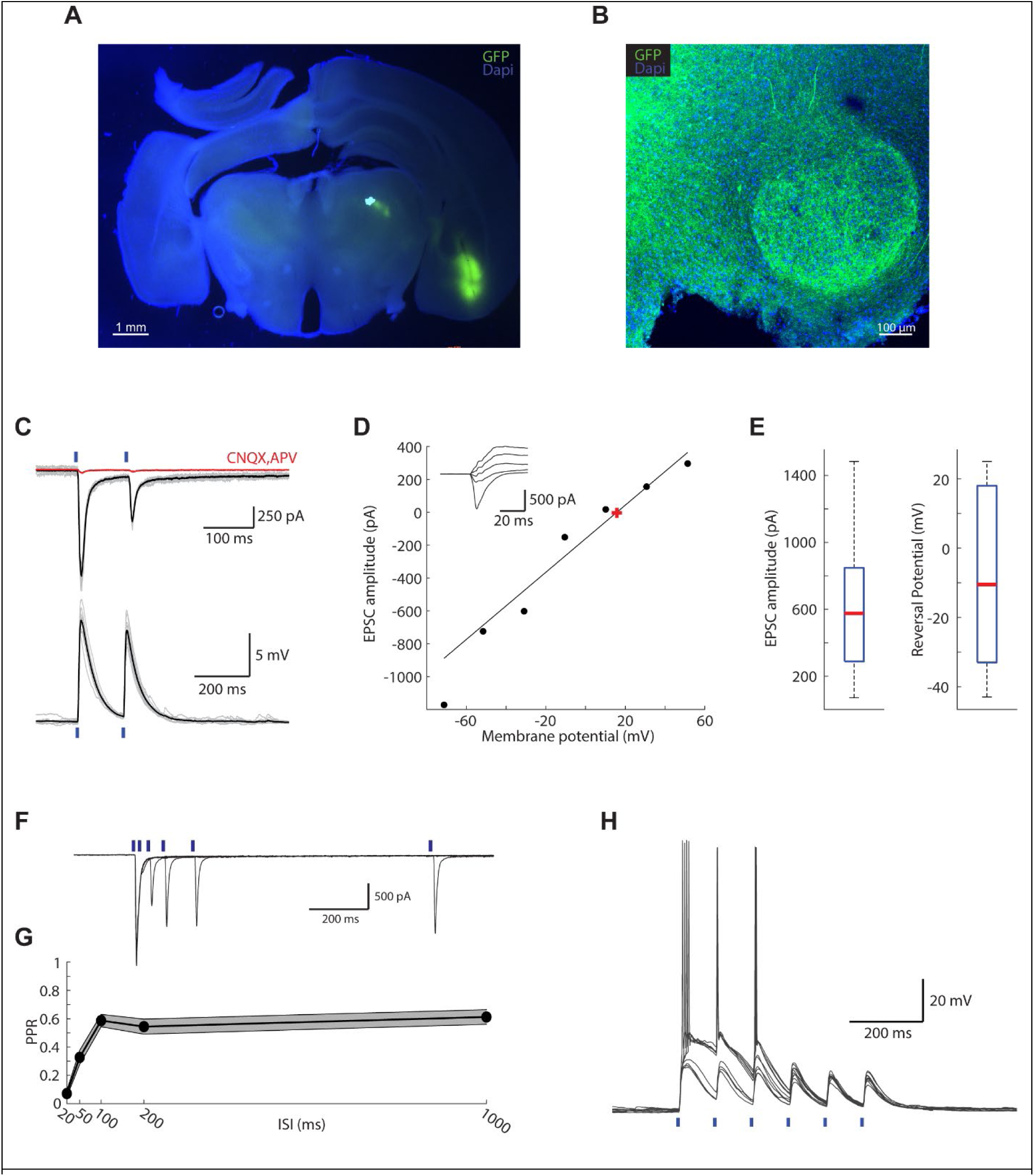
BLA inputs into NLOT. **A.** A coronal section showing expression of ChR2-GFP in the BLA. **B.** A confocal image showing the spread of BLA axons within the NLOT. **C.** Example BLA-evoked EPSCs (top) and EPSPs (bottom). Gray traces show individual trials and black traces show their average. Red trace shows the EPSC obtained in the presence of CNQX and APV. Blue ticks mark the timing of light stimulation. **D.** Voltage dependence of BLA-evoked EPSCs from one example cell. Black line is a linear fit. Red cross denotes reversal potential. **E.** Box plots showing the distributions of amplitudes (n=25 cells) and reversal potentials (n=7 cells) of BLA-evoked EPSCs. **F-G.** Short-term synaptic plasticity of BLA inputs into NLOT. **F.** Example repeated stimulation with variable intervals. **G.** Average paired-pulse ratio as a function of inter-pulse interval. Shaded area shows the mean±SEM (n=15 cells). **H.** Example BLA-evoked action potential bursts. 5 repetitions of the same stimulation are shown.

In summary, NLOT neurons are most efficiently activated by PC and BLA inputs. These inputs show paired-pulse depression which limits the time of activation with repeated stimulus trains.

## Discussion

We combined virus-based tracing methods with acute slice electrophysiology to describe the biophysical properties of NLOT neurons and their synaptic connectivity. We describe the physiological properties of synaptic inputs into the NLOT from olfactory brain regions and from the BLA.

We find that NLOT neurons are bursters. When injected with minimal current steps, or activated synaptically, they respond with a single short burst of action potentials. These bursts are dependent on a calcium current and are typically brief (100-200 ms) with action potentials occurring at a high rate of over 30 Hz. Ca^2+^-current dependent bursts are common in various neuron types including several types of cortical pyramidal neurons (Connors and Gutnick, 1990; Markram et al., 2004). Such bursting has been suggested to be transmitted across synapses with much greater reliability as the high rate of action potentials ensures temporal summation of postsynaptic potentials and may be specifically efficient in facilitating synapses (Lisman, 1997; Izhikevich, 2006). However, bursting also means that the neuronal output may act in a binary fashion rather than a continuous firing rate value and therefore may have a lower encoding capacity. Bursting of NLOT neurons may therefore convey a low-dimensional signal that is of high importance and needs to efficiently activate its postsynaptic targets. The firing properties of NLOT neurons seem distinct from the properties of other amygdalar excitatory neurons. These have been shown to be variable both molecularly and biophysically, ranging in their firing properties from adapting to continuous firing neurons (Rainnie et al., 1993; Faber et al., 2001; Lopez De Armentia and Sah, 2004; O’Leary et al., 2020). We find that layer 2 neurons are quite homogeneously bursting neurons. The distinct properties of NLOT neurons as compared to other amygdalar neurons is in line with their unique gene expression profile (Hochgerner et al., 2023).

Using virus-based tracing methods we find that the main input sources as well as output targets of NLOT are olfactory and amygdalar brain regions. Some of the connectivity that we describe here has been previously shown using other techniques (Price, 1973; Scalia and Winans, 1975; Krettek and Price, 1978; Luskin and Price, 1983; Santiago and Shammah-Lagnado, 2004; Krabbe et al., 2019). Olfactory brain regions represent the largest fraction of NLOT inputs and outputs followed by BLA, cortical amygdala nuclei, and internal NLOT connectivity. Additionally, we show here that there are two inputs from brain regions that contain cholinergic neurons – the SI and the NDB, which may explain the dense expression of acetylcholine esterase in the NLOT (Millhouse and Uemura-Sumi, 1985). These may play a role in state-dependent modulation NLOT function. Interestingly, the projections from the NLOT to many brain regions are either bi-hemispheric, or exclusively contralateral. The NLOT projects bilaterally to the more anterior structures (OB, aPC, OT), and contralaterally to the more posterior structures (pPC and BLA). The significance of this interhemispheric projection pattern is yet unclear.

The input from the OB is rather weak and incapable of driving NLOT output by itself, much different than OB input into PC (Franks and Isaacson, 2005, 2006; Wiegand et al., 2011). On the other hand, inputs from olfactory cortical regions (PC and TT/AON) are powerful and can drive NLOT bursting. This suggests that NLOT activity primarily reflects highly processed olfactory information (Tanisumi et al., 2021). Interestingly, similar findings were described for olfactory inputs into the medial amygdala (Keshavarzi et al., 2015). There, direct input from the accessory OB is much weaker (and located distally on the dendrite) compared to the more processed cortical amygdala input. What may be the impact of the OB input that is located so distally on the dendrite? The potential effects of distal inputs on neuronal output have been studied both experimentally and theoretically (Bernander et al., 1994; Häusser, 2001). Two findings seem relevant to the NLOT. First, distal inputs may be effective in modifying spike output when they coincide with proximal inputs. This coincidence is particularly effective in activating dendritic voltage gated channels and eliciting dendritic action potentials (Larkum et al., 1999). Indeed, we find that when NLOT neurons are depolarized, OB inputs may elicit regenerative responses that are most probably dendritic spikes. Second, distal inputs may induce plastic changes of proximal input responses. Such interaction has been reported in hippocampal CA1 neurons (Dudman et al., 2007).

What can we learn from the synaptic properties of NLOT about its function? We find that the major sources of glutamatergic input into NLOT are from olfactory brain regions and from the BLA. Based on this integration of sensory and emotional signals, it is parsimonious to hypothesize that NLOT is involved in assigning emotional value to specific smells. The fact that NLOT also projects back to olfactory regions, including the OB, may suggest that it may also play a role in modifying olfactory perception based on emotional state. Recording the activity of NLOT neurons during carefully designed behavioral paradigms may shed more light on this hypothesis.

